# Novel estrogen replacement combination therapy including the investigational drug davunetide

**DOI:** 10.64898/2026.05.20.726476

**Authors:** Liri Sophia Guz, Artur Galushkin, Illana Gozes

## Abstract

Estrogen is an essential hormone that critically impacts bodily and brain functions, supporting learning, memory, and motor activities. A decrease in estrogen levels is associated with cognitive decline and motor dysfunction, such as muscle weakness. While conventional hormone replacement treatments (HRT) exist, those have limitations and potentially severe side effects. NAP (davunetide) is the smallest neuroprotective peptide site of activity-dependent neuroprotective protein (ADNP), a master regulator of cognition, essential for brain formation. It is known that NAP restores ADNP activity in cases of deficiency and it has already shown potential in preventing cognitive impairment, protecting against tauopathy, and improving motor function in various animal models and in clinical trials. Based on the dynamic regulation of ADNP by the estrous cycle and its involvement in steroidogenic pathways, we hypothesize that NAP may restore ADNP activity and thus serve as an alternative to conventional hormonal treatments. To test this, 3-month-old female ICR mice underwent bilateral ovariectomy (OVX) or Sham surgery and received daily intranasal administration of NAP, estrogen, or vehicle. Results showed a significant reduction in weight-normalized forelimb grip strength in the OVX model. Daily administration of NAP or estrogen resulted in intermediate grip strength levels that did not statistically differ from either the Sham control or untreated OVX groups. Interestingly, grip strength was the only test that yielded significant results, and no significant differences were observed in the Novel Object Recognition (NOR) test or computed tomography (CT) scans. These findings suggest that NAP may effectively prevent the loss of physical force production typically seen following ovarian hormone depletion, presenting a viable, non-hormonal candidate strategy for managing musculoskeletal symptoms. We hypothesize that the lack of significance in other parameters was due to soy-derived phytoestrogens in the diet, which may have exerted a systemic estrogenic effect that masked the expected physiological phenotypes typically observed in OVX models. Future replication using phytoestrogen-deficient food is required to isolate the specific neuroprotective and musculoskeletal effects of NAP from dietary influence and clarify the broader therapeutic benefits of NAP.

## 2. Introduction

Estrogen is an essential hormone that critically impacts bodily and brain functions. It supports learning and memory processes, influences mood, and is important for maintaining bone strength, cardiovascular health, sexual health, and many other functions (Casey, 2017). A decline in this hormone’s levels, whether due to aging during menopause or other physiological conditions, is associated with an increased risk of tauopathy in the brain (Russell et al., 2019) and can lead to cognitive decline, memory problems, decreased motivation, impaired learning abilities, and motor dysfunction, such as muscle weakness or osteoporosis.

Currently, conventional hormone replacement treatments (HRT) exist, but these treatments have limitations and potentially severe side effects, such as an increased risk of blood clots and cancer development (NHS, 2023). Additionally, there are medical conditions where these treatments are not feasible or recommended. For example, women with a history of breast cancer, especially estrogen-dependent breast cancer, may face an increased risk of cancer recurrence with estrogen therapy. Other examples include men with a history of prostate cancer or individuals of either sex with a history of blood clotting disorders or hypertension, as hormonal treatments may exacerbate these conditions (NHS, 2023). Moreover, some individuals choose to avoid hormonal replacement therapy due to side effects such as bloating, fluid retention, breast tenderness, and reduced fertility in men. While these side effects are not life-threatening, they can negatively affect quality of life and should be considered when initiating treatment (NHS, 2023).

To investigate the mechanisms underlying these estrogen-related declines and to test potential therapeutic alternatives to HRT, researchers often utilize the ovariectomized (OVX) animal model. The surgical procedure bilaterally removes the ovaries, which results in a rapid and significant reduction in endogenous estrogen levels, effectively mimicking the physiological state of menopause in humans (Souza et al., 2018).

In mice, the OVX model reliably reproduces many of the clinical symptoms described in humans. Phenotypically, these mice exhibit a decrease in bone mineral density like human osteoporosis (Xu et al., 2022). This decline is frequently accompanied by a measurable cognitive impairment such as in the Novel Object Recognition test and the Morris water maze test (Tao et al., 2020), and a decrease in the muscle force generation of OVX mice, measured for example in the grip test analysis (Kitajima and Ono, 2016).

Our previous studies identified NAP (NAPVSIPQ single amino acid letter code, also known as davunetide, AL-108, CP201 and sometimes referred to as NAP peptide) as the smallest neuroprotective peptide site of activity-dependent neuroprotective protein (ADNP) (Bassan et al., 1999), a master regulator of cognition, essential for brain formation (Galushkin & Gozes, 2025; Pinhasov et al., 2003; Vulih-Shultzman et al., 2007). It is known that NAP restores ADNP activity in cases of deficiency and it has already shown potential in preventing cognitive impairment, protecting against tauopathy, and improving motor function in various models (Karmon et al., 2022; Hacohen-Kleiman et al., 2019), both in animals and humans, such as in studies on progressive supranuclear palsy (Gozes et al., 2023; Gozes et al., 2026) and pre-Alzheimer’s disease conditions (Gozes et al., 2024; Gozes et al., 2025). NAP and pipeline products protect nerve cells by linking with microtubule end binding proteins (EB1/EB3), through the embedded EB1/EB3 interacting SxIP motif (NAPV**SIP**Q, Ser-Ile-Pro), thus enhancing microtubule dynamics and Tau-microtubule interaction, protecting the synapse. NAP further enhances ADNP-EB1/EB3, interactions, ameliorating ADNP deficits (Gozes & Shazman, 2023; Hacohen-Kleiman et al., 2018; Ivashko-Pachima & Gozes, 2021).

While originally characterized by its potent neuroprotective properties and its ability to stabilize microtubules (Ivashko-Pachima and Gozes, 2021), research has identified a significant sexual dimorphism in the expression of ADNP. ADNP expression is not static, rather, it exhibits significant sexual dimorphism and fluctuations in the arcuate nucleus of the mouse hypothalamus with the female estrus cycle. Specifically, ADNP-like immunoreactivity peaks during proestrus, the phase characterized by maximal levels of estradiol and progesterone necessary to trigger the preovulatory luteinizing hormone (LH) surge (Furman et al., 2004).

Moreover, the importance of ADNP extends beyond its role as a passive marker of hormonal shifts as it appears to be a regulator of the very pathways that produce these hormones-transcriptomic analyses of ADNP-mutant lymphoblastoid cells have revealed significant alterations in the expression of genes belonging to the steroid-biosynthesis pathway. Gene-set enrichment analysis (GSEA) of these mutants identified “steroid-biosynthesis” as a top hit among significantly over-represented KEGG pathways (P < 0.05) (Grigg et al., 2020).

Our recent findings indicated ADNP/sex regulation of the unfolded protein response (UPR), a stress response that is activated upon detection of excess unfolded or misfolded proteins within the lumen of the endoplasmic reticulum (ER), with decreased levels of estrogen or testosterone in females or males leading to an increase in ER stress due to dysregulation of the UPR, thereby contributing to conditions, diseases or disorders associated with low sex hormone (Shapira et al., 2025). Importantly, NAP treatment protects against ADNP deficiency (mutation) associated with UPR function (Shapira et al., 2025). Interestingly, this regulation exhibits distinct sexual dimorphism and while UPR dysregulation is more pronounced in males, females display a stronger association with mitochondrial regulation.

Thus, we hypothesize that NAP (the investigational drug, davunetide) will constitute a novel hormone replacement therapy. Testing our hypothesis in the OVX model indeed showed a significant decrease in grip strength in the model that was ameliorated by either NAP or estrogen treatment.

## 3. Methods and Materials

### 3.1. Animals

All procedures involving animals have been approved by the Animal Care and Use Committee of Tel Aviv University and the Israeli Ministry of Health. A total of 24 3-month-old female ICR mice were ordered from Harlan and were used in this study. All mice were group-housed, kept on a 12 h light/dark cycle, and had free access to rodent chow and water at the Glasberg Animal Tower of the Gray Faculty of Medical and Health Sciences at Tel Aviv University.

Mice were randomly allocated into four experimental groups: Control (Sham-operated, receiving Vehicle), OVX (Vehicle), OVX + NAP, and OVX + Estrogen. Treatments were administered via daily intranasal administration (5 µL per mouse) five days per week for the duration of the study. The NAP group received a dosage of 0.5 µg/5 µL vehicle per day, while the estrogen group received a final concentration of 1 µg per 5 µL dose. Control and OVX-only groups received an equivalent volume of vehicle (1×DD) (Alcalay et al., 2004).

### 3.2. Dietary Composition and Phytoestrogen Analysis

During the experimental period, mice were maintained on the Altromin 1318 standardized diet. The dietary formulation contains soybean meal, soy husks, and soybean oil. Given that soy-derived isoflavone content can fluctuate significantly based on environmental and harvest conditions, a sample batch-specific analysis was utilized to characterize the phytoestrogen profile of the diet. The analyzed concentrations for the specific representative batch used were 0.362 mg/g Genistein, 0.298 mg/g Daidzein, and 0.068 mg/g Glycitein, resulting in a total isoflavone content of 0.727 mg/g.

### 3.3. Experimental Design and Timeline

The experimental design and timeline are outlined in Figure 1.

**Figure 1.**
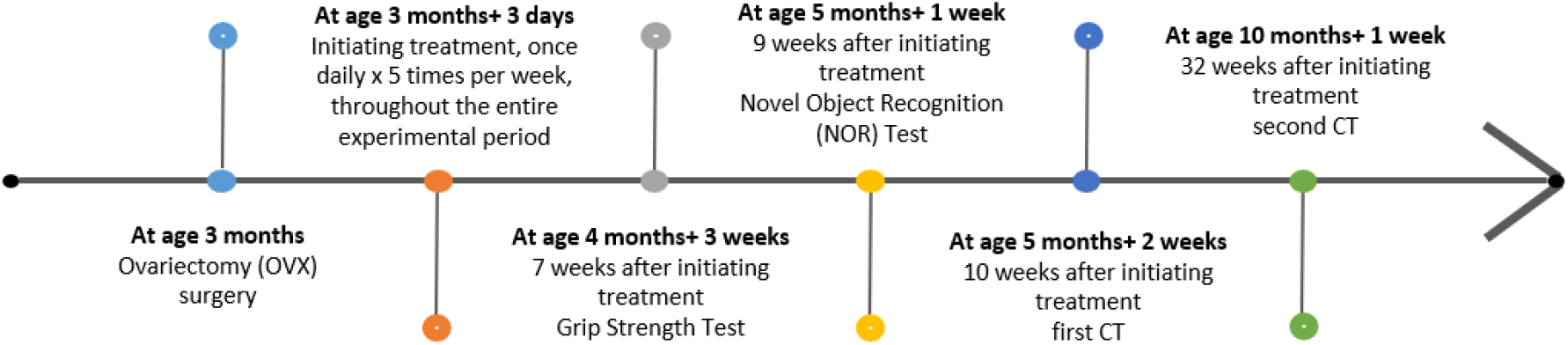
Experimental Design and Timeline. Female mice (n= 24) underwent either bilateral ovariectomy (OVX) or Sham surgery at 3 months of age. Following a few days of recovery, mice were randomly assigned to four experimental groups: Sham + Vehicle, OVX + Vehicle, OVX + NAP, OVX + estrogen. Daily intranasal administration was maintained for the whole duration of the study. Functional and behavioral tests including Grip Strength, Novel Object Recognition (NOR) and CT imaging were conducted.

### 3.4. Ovariectomy (OVX) surgery

Surgical ovariectomy was performed under general anesthesia induced by an intraperitoneal injection of a Ketamine (100 mg/kg) and Midazolam (1 mg/kg) cocktail. To ensure animal welfare and minimize distress, pre- and post-operative analgesia was administered using Carprofen (5 mg/kg), provided subcutaneously as needed based on clinical assessment.

Following the induction of anesthesia, a midline dorsal incision was made, and the ovaries were bilaterally localized, ligated, and excised. Sham-operated mice underwent the same surgical exposure without ovary removal to serve as a procedural control. All animals were allowed a five-day recovery period with close monitoring before the initiation of intranasal treatments.

### 3.5. Behavioral Tests

All behavioral tests were conducted during the light phase of the circadian cycle in a dedicated, sound-attenuated room. Mice were acclimated to the testing environment for at least 60 minutes prior to the start of each session. All equipment was thoroughly cleaned with a disinfectant solution (Virosol) between trials to eliminate olfactory cues.

#### 3.5.1. Grip Strength Test

Forelimb grip strength was quantified using a digital Grip Strength Meter (Ugo Basile, Model 47200). Mice were held by the tail and lowered over the grip bar, ensuring the torso remained horizontal. Once the mouse firmly grasped the bar with its two forepaws, it was gently and steadily pulled backward until enough tension was created between the mouse and the bar and was held there until the grip was released. The peak pull force was recorded in grams force (gf) by the digital force transducer.

To ensure accuracy and minimize variability, the gauge was reset to zero before each trial. Each animal underwent five consecutive trials in a single session. The three highest recorded values were averaged to determine the final grip strength score for each mouse. This “top three” averaging method was employed to account for fluctuations in mouse engagement or improper attachment during individual trials.

Numerous studies to date have reported that muscle strength positively correlates with body weight, and studies that use muscle strength as an outcome variable usually adopt a method of normalizing this variable by body mass. Additionally, grip strength is correlated with muscle mass. Therefore, absolute grip strength values (gf) were normalized to individual body weight (g), yielding relative grip strength expressed in grams force per gram of body weight (gf/g) (Chun et al., 2019).

#### 3.5.2. Novel Object Recognition (NOR) Test

The test is based on visual discrimination between two different objects in an arena of 50 cm x 50 cm, conducted over three consecutive days. The first day is the habituation day, where each mouse is placed in the empty testing arena for 5 min. This part is also utilized as an open field test. On the second day (the learning phase), each mouse is placed in the same arena with 2 identical objects and allowed to explore them for 5 min, while the time spent sniffing/ touching each object is recorded. On the third and last day (the testing phase), each mouse is placed in the same arena for 5 min each, with one familiar object removed and replaced with a novel object. The time spent exploring each object was recorded and a Discrimination Index (DI) was calculated to determine the preference for the novel object.

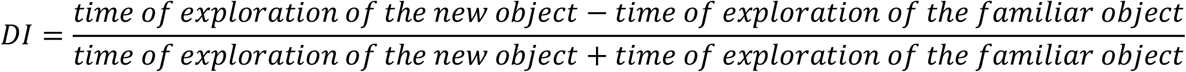

A minimum exploration threshold was established-any mouse with a total exploration time of less than 30s during either the second or the third day was excluded from the final statistical analysis.

#### 3.5.3. Computed tomography (CT) Imaging

The mice underwent X-ray computed tomography (CT) using the VECTor4 system from MILabs. Mice were anesthetized with an intraperitoneal injection of a Ketamine (100 mg/kg) and Midazolam (1 mg/kg) cocktail. Quantitative analysis was focused on the femur bone.

### 3.6. Statistical Analysis

Statistical analysis was performed using GraphPad Prism version 10.6.1 software.

## 4. Results

### 4.1. Grip Strength Assessment

To evaluate the impact of OVX and the potential protective effects of NAP, forelimb grip strength was measured seven weeks after initiating treatment, (mice aged 4 months and 3 weeks) and normalized to total body weight.

As shown in Figure 2, OVX induced a minor but statistically significant reduction in forelimb grip strength compared to Sham-operated controls (P = 0.0184). Daily intranasal administration of either NAP or estrogen resulted in an increase in the weight-normalized grip strength compared to the untreated OVX group that did not statistically differ from either the Sham control or the untreated OVX groups.

**Figure 2.**
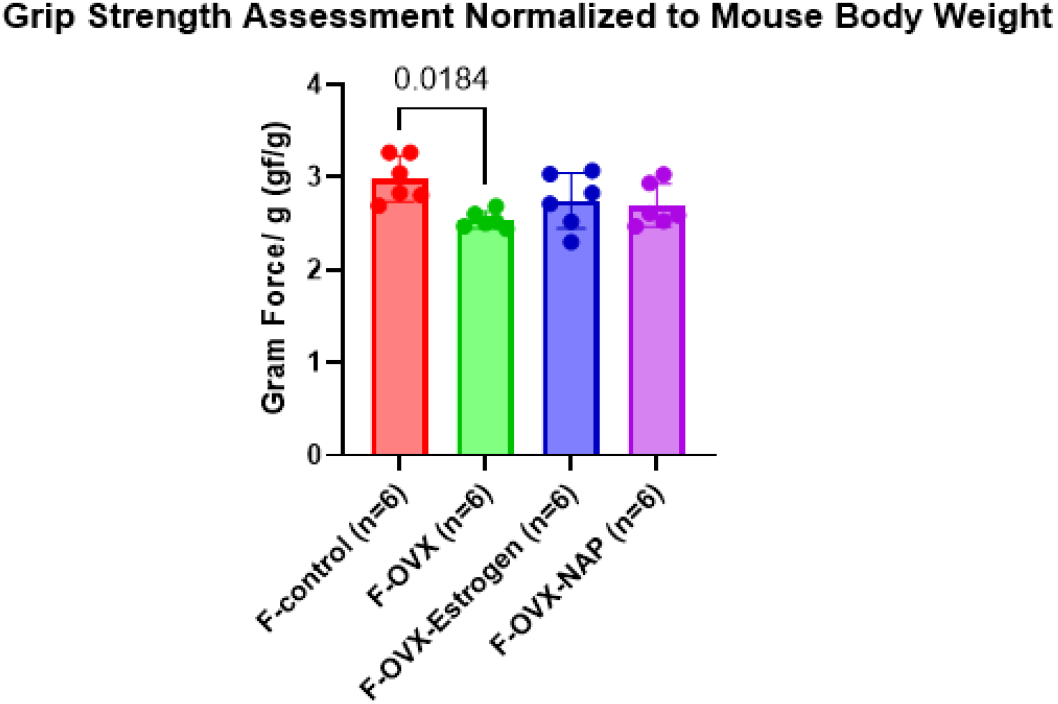
Weight-Normalized Forelimb Grip Strength Assessment at 7 Weeks Post-Treatment Initiation. Forelimb grip strength was measured 7 weeks after initiating treatment and normalized to individual mouse body weight, expressed as grams force per gram of body weight (gf/g, n=6 per group). Ovariectomy (F-OVX) induced a statistically significant reduction in normalized grip strength compared to the Sham-operated control group (F-control, P = 0.0184). Daily intranasal administration of either estrogen (F-OVX-Estrogen) or NAP (F-OVX-NAP) resulted in grip strength levels that did not statistically differ from either the control or untreated OVX groups. Data passed the Shapiro-Wilk normality test and differences between groups were analyzed using a one-way ANOVA followed by the Holm-Sidak post-hoc test for multiple comparisons.

### 4.2. Novel Object Recognition (NOR) Assessment

To evaluate the impact of OVX and the potential protective effects of NAP, the Novel Object Recognition (NOR) test was performed 9 weeks after initiating treatment, when the mice were 5 months and 1 week old.

On the second day of the NOR assessment, object side preference was evaluated during the learning phase. A two-tailed t-test confirmed that there were no significant differences in the duration spent exploring the identical objects A1 vs. A2 across all experimental groups (Figure 3. A).

**Figure 3.**
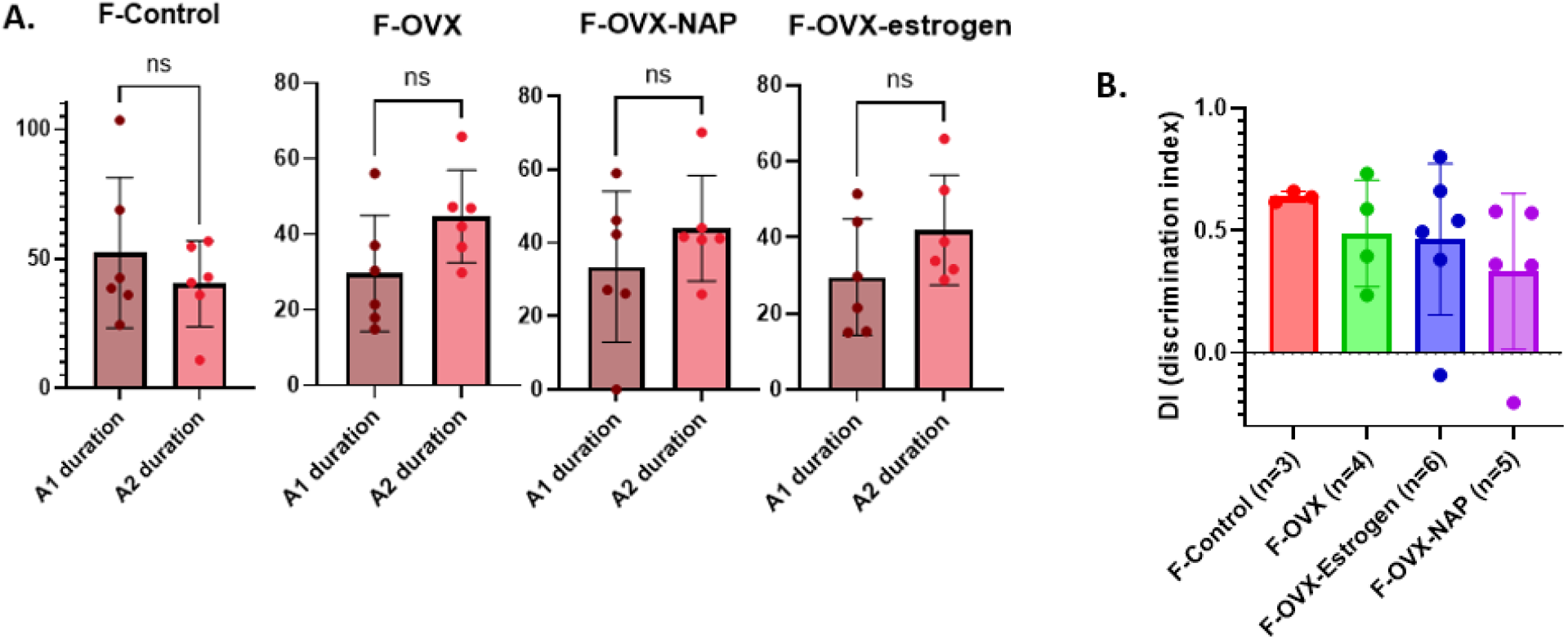
Object Recognition Assessment at 9 Weeks Post-Treatment Initiation. **(A)** Duration (s) spent exploring two identical objects-A1 and A2 during the learning phase was recorded for each mouse. Results show no significant differences between the exploration times for objects A1 and A2 in any group, confirming the lack of innate side bias prior to the testing phase of the NOR test. P calculated by Mann-Whitney U Test. **(B)** Recognition ability was evaluated using the Discrimination Index (DI). The DI was calculated for each experimental group: Control (F-Control, n=3), Ovariectomized (F-OVX, n=4), and OVX treated with NAP (F-OVX-NAP, n=5) or Estrogen (F-OVX-Estrogen, n=6). Results show no significant differences between the groups, indicating that at 9 weeks’ post-treatment, all groups exhibited similar levels of preference for the novel object compared to the familiar object. Data passed the Shapiro-Wilk normality test. Differences between groups were analyzed using a one-way ANOVA followed by the Holm-Sidak post-hoc test for multiple comparisons.

On the third day (the testing phase), recognition memory was evaluated using the Discrimination Index (DI). Statistical analysis showed that all groups exhibited a similar preference for the novel object, and no significant differences were observed between all experimental groups (P > 0.05) (Figure 3. B).

### 4.3. CT Imaging Assessment

To evaluate the impact of OVX and the potential protective effects of NAP, CT analysis of the femur bone was conducted at two different time points including 10 (Figure 4) and 32 (Figure 5) weeks post-treatment initiation.

**Figure 4.**
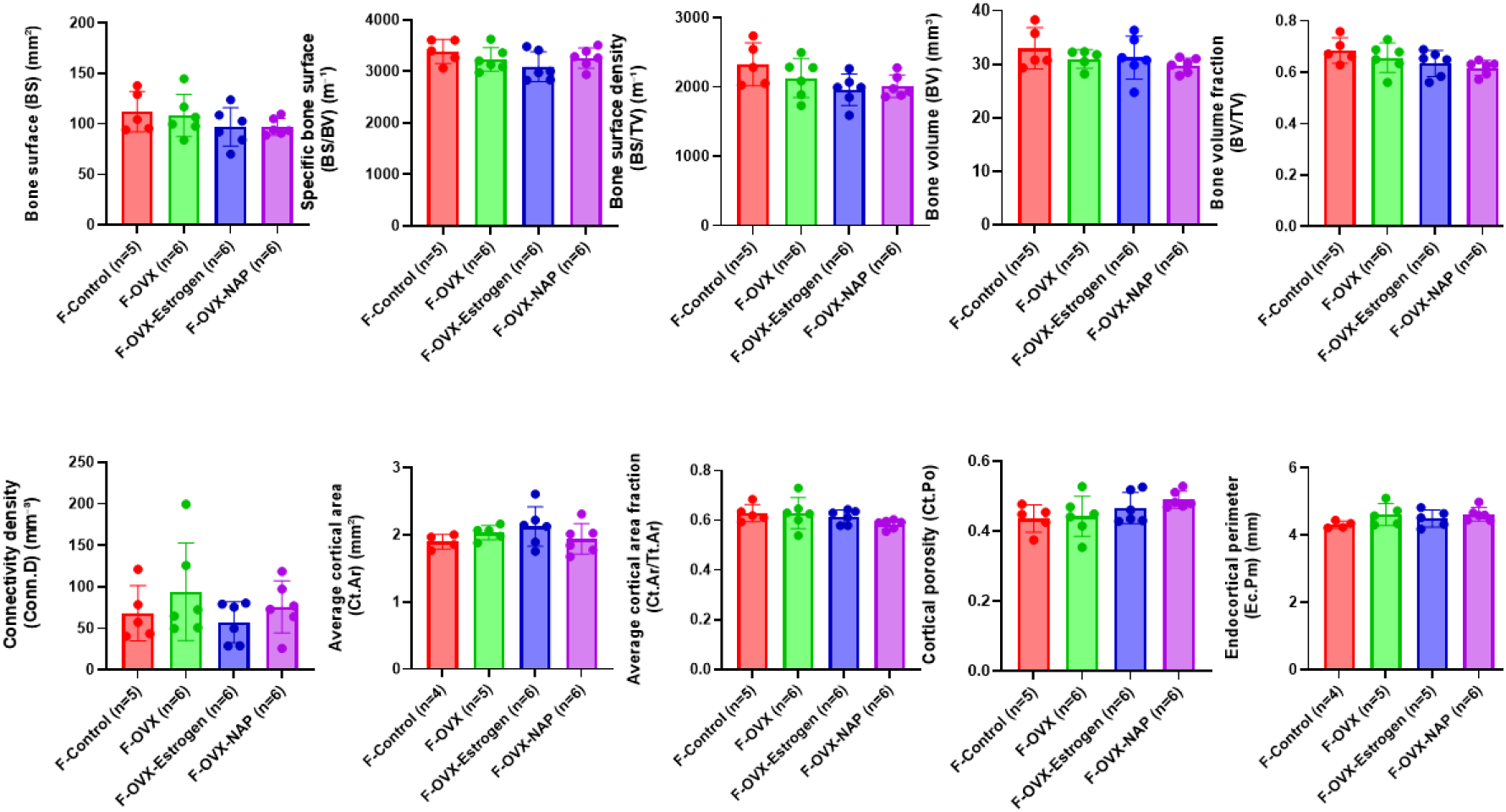
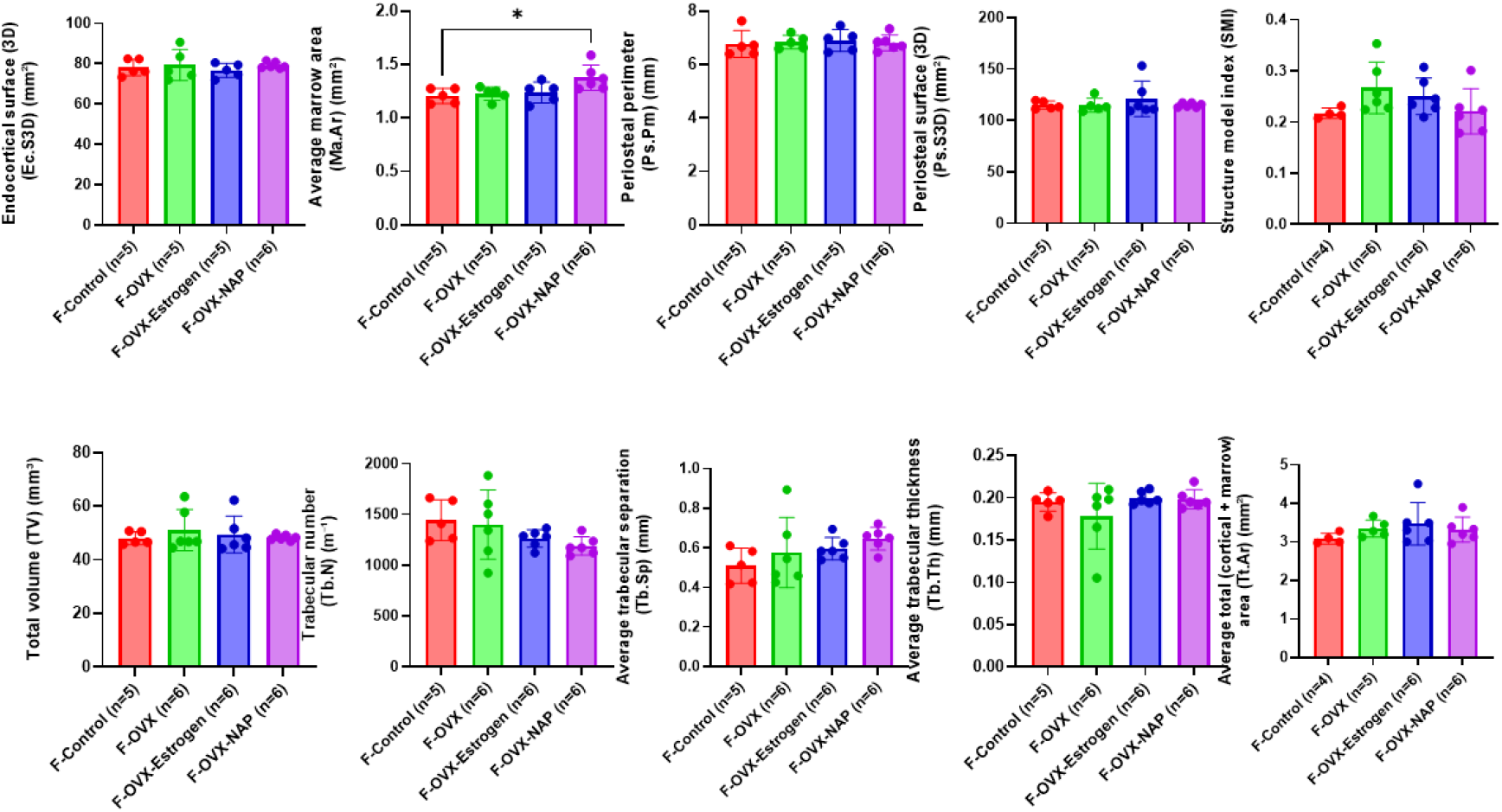
CT Imaging Assessment 10 Weeks Post-Treatment Initiation. Structural bone integrity was assessed 10 weeks after the initiation of treatment. Multiple cortical and trabecular parameters were analyzed for Control (F-Control), Ovariectomized (F-OVX), and OVX treated with NAP (F-OVX-NAP) or Estrogen (F-OVX-Estrogen). Results show no significant differences between any of the experimental groups, indicating that at 10 weeks’ post-treatment initiation, skeletal architecture remained comparable across all conditions. Data passed the Shapiro-Wilk normality test. Differences between groups were analyzed using a one-way ANOVA followed by the Holm-Sidak post-hoc test for multiple comparisons.

**Figure 5.**
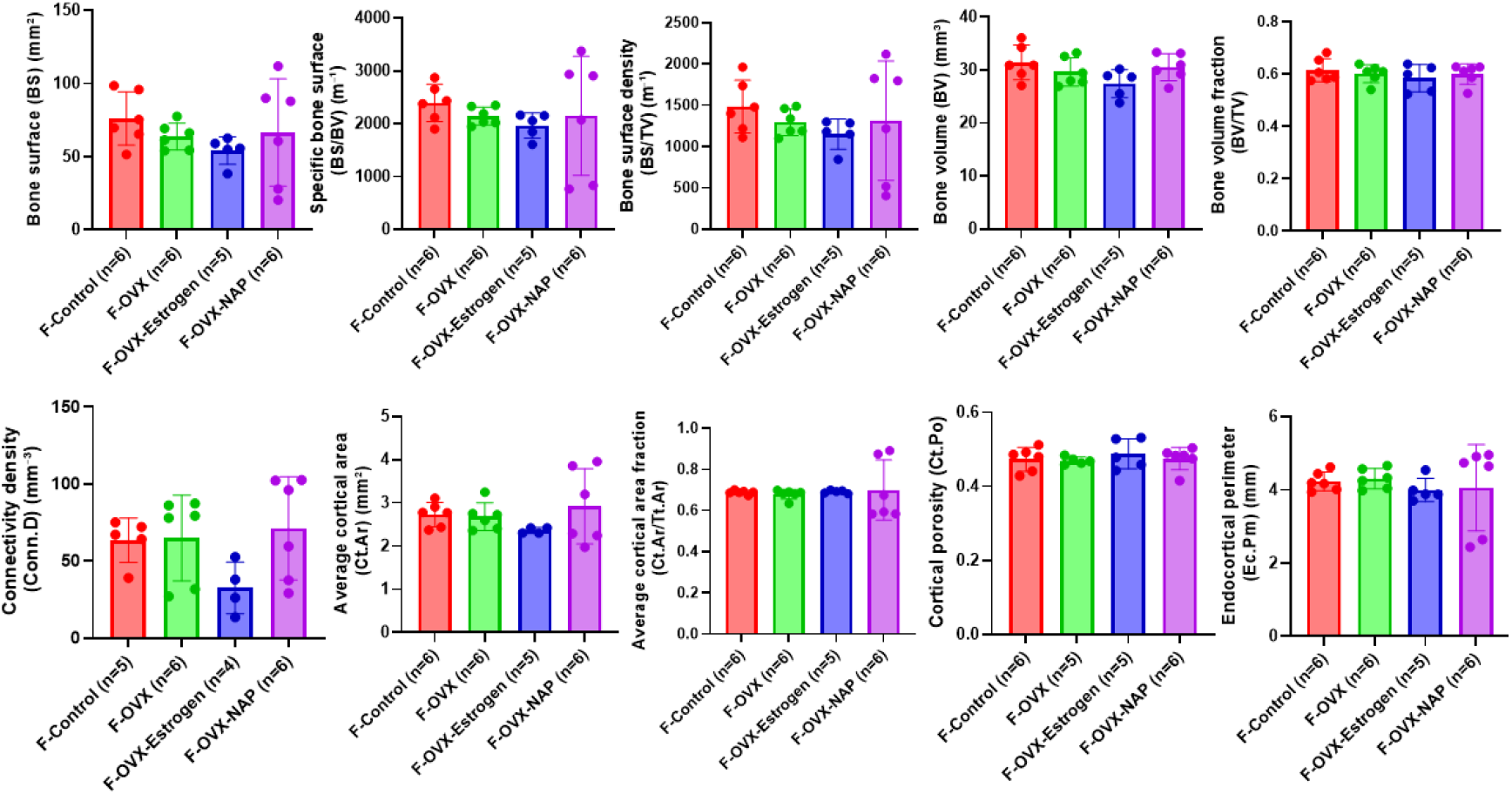
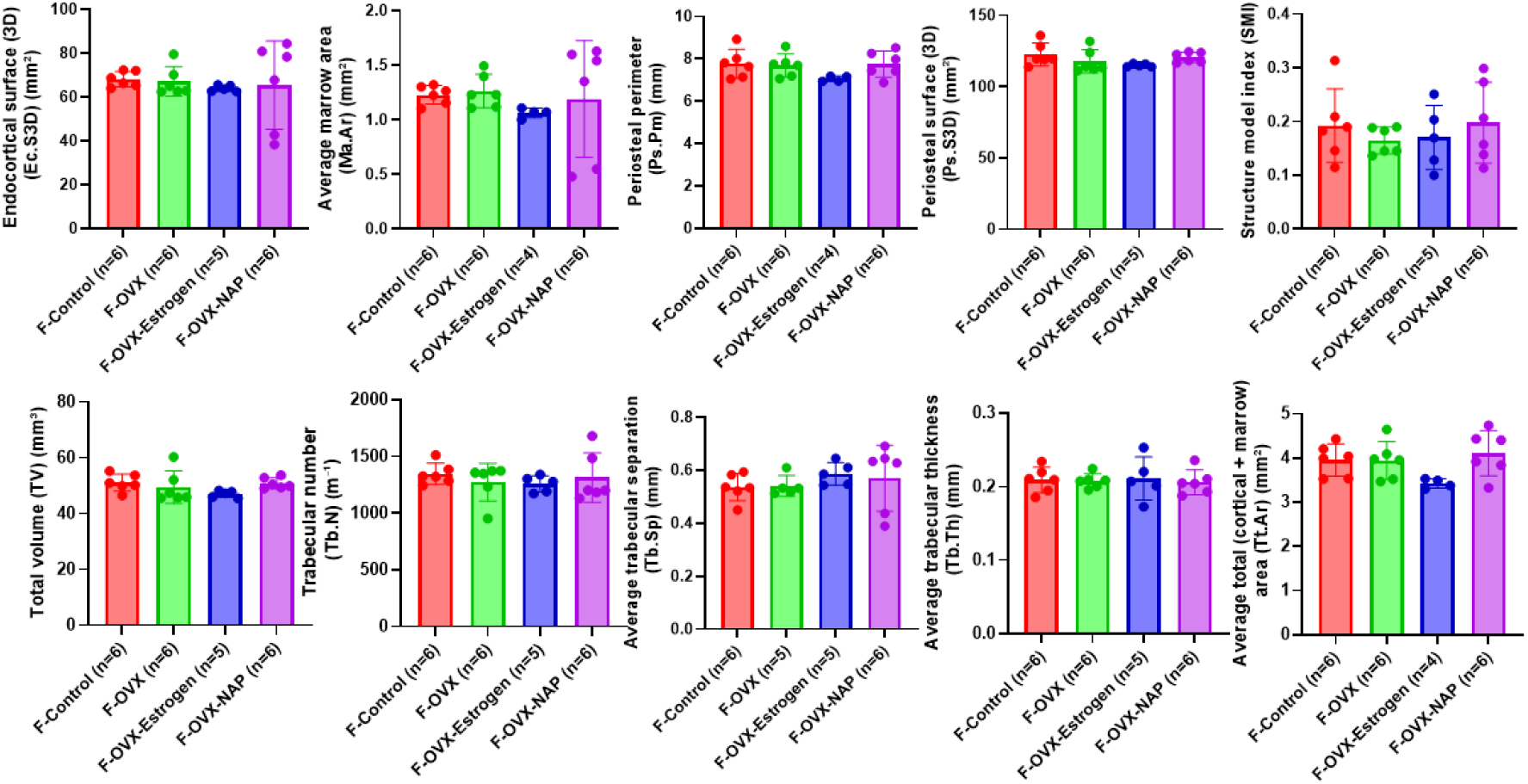
CT Imaging Assessment 32 Weeks Post-Treatment Initiation. Structural bone integrity was assessed 32 weeks after the initiation of treatment. Multiple cortical and trabecular parameters were analyzed for Control (F-Control), Ovariectomized (F-OVX), and OVX treated with NAP (F-OVX-NAP) or Estrogen (F-OVX-Estrogen). Results show no significant differences between any of the experimental groups, indicating that at 32 weeks post-treatment initiation, skeletal architecture remained comparable across all conditions. Data passed the Shapiro-Wilk normality test. Differences between groups were analyzed using a one-way ANOVA followed by the Holm-Sidak post-hoc test for multiple comparisons.

At 10 weeks post-initiation, no significant differences (P > 0.05) were observed between the experimental groups in the evaluated parameters. Moreover, despite the extended duration of the study and the prolonged state of estrogen deficiency in the OVX groups, no significant differences were shown between the experimental groups in the same parameters when re-evaluated at 32 weeks post-initiation (P > 0.05).

## 5. Discussion

To accurately assess grip strength, we evaluated force production relative to body weight, directly controlling for individual mass variations. Numerous studies to date have reported that muscle strength positively correlates with body weight, and studies that use muscle strength as an outcome variable usually adopt a method of normalizing this variable by body mass (Chun et al., 2019). Utilizing this mathematical adjustment ensures that the significant reduction in relative grip strength observed in the untreated OVX group (Fig. 2) reflects a true decline in functional capacity rather than a passive byproduct of total body weight shifts.

This observed deficit in the OVX group likely involves a combination of altered central nervous system (CNS) regulation and intrinsic, structural changes within the musculoskeletal tissue itself.

This hypothesis is supported by recent connectome-based modelling by Ghaffari et al., 2025. In their study, they tested whether they could computationally predict an individual’s handgrip strength, a key clinical biomarker for frailty, by using their fMRI brain scans. Utilizing connectome-based predictive modeling (CPM), they demonstrated that individual variations in physical frailty, specifically handgrip maximum voluntary contraction (MVIC) strength, can be predicted by a person’s unique brain connectome. The study revealed that an individual’s distinct brain-wiring pattern accurately predicts grip strength, identifying the caudate nucleus as the most critical region for predicting physical force. This suggests that “frailty” might start with changes in the brain’s motor networks before it even shows up as major muscle loss.

Furthermore, while NAP is widely documented to enhance synaptic plasticity and stabilize microtubules within the CNS (Karmon et al., 2022; Hacohen-Kleiman et al., 2019), it also exerts direct protective effects within peripheral muscle tissue itself. Single-cell transcriptomics identified ADNP as a major constituent of the developing human muscle, and its systemic deficiency in *Adnp*^*+/-*^ mice has been shown to cause profound neuromuscular junction (NMJ) disruption. Specifically, evaluation of these NMJs revealed significantly decreased microtubule (tubulin) intensities in *Adnp*^*+/-*^ compared to *Adnp*^*+/+*^ muscles, demonstrating a significant genotype-related structural reduction in males, but not in females. Crucially, significant positive correlations were discovered between behavioral test performance and localized *Adnp* gene expression levels in the muscles of 7-month-old mice. Additionally, local muscle deficiency achieved via CRISPR/Cas9 knockdown of adult mouse gastrocnemius *Adnp* resulted in severe motor dysfunctions that manifested in a distinctly sex-dependent manner. Mechanistically, ADNP was found to regulate both muscle microtubules and myosin light chain (Myl2), and these localized deficits were specifically ameliorated by daily administration of NAP (Kapitansky et al., 2020).

In the context of our study, the protective trend of NAP then may stem from this dual mechanism of action-its established role in stabilizing microtubules and enhancing synaptic plasticity alongside its stabilization of peripheral muscle architecture.

Interestingly, weight-normalized grip strength was the only test that yielded significant results in our study. We hypothesize that the lack of significance in the Novel Object Recognition (NOR) and CT scans may be due to the specific composition of the rodent diet, specifically the presence of soy-derived phytoestrogens. These plant-derived compounds, such as genistein and daidzein, bear a striking structural similarity to 17β-estradiol (E2) (Desmawati & Sulastri, 2019), and function as Selective Estrogen Receptor Modulators (SERMs) (Oseni et al., 2008). These compounds may have exerted a systemic estrogenic effect that was sufficient to partially mask the expected physiological phenotypes typically observed in OVX models, thereby narrowing the measurable therapeutic window for NAP.

To address this potential interference and isolate the specific neuroprotective and musculoskeletal effects of NAP from a possible dietary confounding influence, our next experimental step will involve a complete replication of this study design using a phytoestrogen-deficient diet. By eliminating the background estrogenic activity provided by soy-based components, we aim to clarify whether a “cleaner” hormonal environment reveals broader therapeutic benefits across the cognitive and physiological parameters that remained unchanged in the current study.

## 6. Conclusions

Our data show that NAP (davunetide) effectively prevents the loss of physical force production typically seen following ovarian hormone depletion. The fact that NAP partially mitigated the decline in normalized grip strength to intermediate levels comparable to estrogen presents it as a viable, non-hormonal strategy for managing the musculoskeletal symptoms of menopause.

This is a critical finding for clinical populations where conventional HRT is contraindicated, such as patients with cardiovascular risks or estrogen-dependent oncological histories. Moreover, as for potential treatments in humans, dietary interventions may provide a practical and non-invasive method for managing the physiological deficits caused by estrogen depletion.

By targeting the brain-to-muscle connection rather than a systemic hormonal fix, we may offer a path toward reducing frailty and improving the quality of life for individuals navigating the complexities of hormone-related decline.

## 7. Author Contributions, Acknowledgments, Funding sources

### 7.1. Data availability

Full preclinical may be made available from the corresponding author upon a reasonable request.

### 7.3. Author Contributions

IG initiated, orchestrated, and supervised the study, acquired funding, and coordinated data collection and analyses, as well as the writing of the paper. LSG performed all behavioral tests, statistical analyses, generated the plots, and actively participated in manuscript writing. AG took part in drug administration.

The manuscript has been read and approved by all authors. All authors take full responsibility for all data, figures, and text and approve the content and submission of the study. No related work is under consideration elsewhere. All authors state that all unprocessed data are available, and all figures provide accurate presentations of the original data.

### 7.4. Funding Resources

The original study was partly supported by the Elton laboratory, Drs. Ronith and Armand Stemmer (French Friends of Tel Aviv University), Anne and Alex Cohen, Canadian Friends of Tel Aviv University, AMN Foundation, and ExoNavis Therapeutics. Funders were not involved in data analysis.

### 7.5. Author Disclosures

IG serves as VP Drug Development at Exonavis Therapeutics Ltd. Developing davunetide for the ADNP syndrome (under patent protection, IG inventor). All other authors confirmed that no conflict of interest exists. The manuscript has been read and approved by all authors.

## Acknowledgements

We thank Dr. Eliezer Giladi for his invaluable assistance in the study. We also thank Dr. Lior Bikovski (manager, Mayers Neuro-Behavioral Core Facility, Gray Faculty of Medical & Health Sciences, Tel Aviv University) for his essential assistance with the behavioral testing infrastructure and assessments, as well as Drs. Yael Zilberstein and Sahar Hiram Bab for their help with mouse imaging and CT analysis (Research Infrastructure and Core Facilities, as above). Furthermore, we are grateful to Altromin Spezialfutter GmbH & Co. KG, Im Seelenkamp 20 D-32791 Lage for their kind assistance with mouse food.

## List of Abbreviations

ADNP: Activity-Dependent Neuroprotective Protein
CNS: Central Nervous System
CPM: Connectome-Based Predictive Modeling
CT: Computed Tomography
DI: Discrimination Index
E2: 17β-Estradiol
EB1/EB3: End-Binding Proteins 1 and 3
ER: Endoplasmic Reticulum
fMRI: Functional Magnetic Resonance Imaging
GSEA: Gene-Set Enrichment Analysis
HRT: Hormone Replacement Therapy
LH: Luteinizing Hormone
MVIC: Maximum Voluntary Isometric Contraction
NMJ: Neuromuscular Junction
NOR: Novel Object Recognition
OVX: Ovariectomy / Ovariectomized
SERM: Selective Estrogen Receptor Modulator
UPR: Unfolded Protein Response

## References

Bassan, M., Zamostiano, R., Davidson, A., Pinhasov, A., Giladi, E., Perl, O., Bassan, H., Blat, C., Gibney, G., Glazner, G., Brenneman, D. E., & Gozes, I. (1999). Complete sequence of a novel protein containing a Femtomolar-Activity-Dependent neuroprotective peptide. Journal of Neurochemistry, 72(3), 1283–1293. 10.1046/j.1471-4159.1999.0721283.x

Chun, S., Kim, W., & Choi, K. H. (2019). Comparison between grip strength and grip strength divided by body weight in their relationship with metabolic syndrome and quality of life in the elderly. PLoS ONE, 14(9), e0222040. 10.1371/journal.pone.0222040

Desmawati, D., & Sulastri, D. (2019). Phytoestrogens and their health effect. Open Access Macedonian Journal of Medical Sciences, 7(3), 495–499. 10.3889/oamjms.2019.086

Furman, S., Hill, J. M., Vulih, I., Zaltzman, R., Hauser, J. M., Brenneman, D. E., & Gozes, I. (2004). Sexual dimorphism of activity-dependent neuroprotective protein in the mouse arcuate nucleus. Neuroscience Letters, 373(1), 73–78. 10.1016/j.neulet.2004.09.077

Galushkin, A., & Gozes, I. (2025). Intranasal NAP (Davunetide): Neuroprotection and circadian rhythmicity. Advanced Drug Delivery Reviews, 220, 115573. 10.1016/j.addr.2025.115573

Ghaffari, A., Abouzaki, M., Romero, Y., Sun, A., Seitz, A., Langley, J., Bennett, I. J., & Hu, X. (2025). Connectome-based predictive modeling of grip strength: a marker of physical frailty. Frontiers in Neuroscience, 19, 1697908. 10.3389/fnins.2025.1697908

Gozes, I., Blatt, J., & Guz, L. S. (2025). Systemic davunetide provides sex-specific neuroprotection during Coronary Artery Bypass Grafting (CABG). Translational Psychiatry, 15(1), 454. 10.1038/s41398-025-03675-y

Gozes, I., Blatt, J., & Lobyntseva, A. (2024). Davunetide sex-dependently boosts memory in prodromal Alzheimer’s disease. Translational Psychiatry, 14(1), 412. 10.1038/s41398-024-03118-0

Gozes, I., Shapira, G., Lobyntseva, A., & Shomron, N. (2023). Unexpected gender differences in progressive supranuclear palsy reveal efficacy for davunetide in women. Translational Psychiatry, 13(1), 319. 10.1038/s41398-023-02618-9

Gozes, I., & Shazman, S. (2023). A novel davunetide (NAPVSIPQ Q to NAPVSIPQ E) point mutation in activity-dependent neuroprotective protein (ADNP) causes a mild developmental syndrome. European Journal of Neuroscience, 58(2), 2641–2652. 10.1111/ejn.15920

Grigg, I., Ivashko-Pachima, Y., Hait, T. A., Korenková, V., Touloumi, O., Lagoudaki, R., Van Dijck, A., Marusic, Z., Anicic, M., Vukovic, J., Kooy, R. F., Grigoriadis, N., & Gozes, I. (2020). Tauopathy in the young autistic brain: novel biomarker and therapeutic target. Translational Psychiatry, 10(1), 228. 10.1038/s41398-020-00904-4

Hacohen-Kleiman, G., Sragovich, S., Karmon, G., Gao, A. Y. L., Grigg, I., Pasmanik-Chor, M., Le, A., Korenková, V., McKinney, R. A., & Gozes, I. (2018). Activity-dependent neuroprotective protein deficiency models synaptic and developmental phenotypes of autism-like syndrome. Journal of Clinical Investigation, 128(11), 4956–4969. 10.1172/jci98199

Hacohen-Kleiman, G., Yizhar-Barnea, O., Touloumi, O., Lagoudaki, R., Avraham, K. B., Grigoriadis, N., & Gozes, I. (2019). Atypical auditory brainstem response and protein expression aberrations related to ASD and hearing loss in the ADNP haploinsufficient mouse brain. Neurochemical Research, 44(6), 1494–1507. 10.1007/s11064-019-02723-6

Hadar, A., Kapitansky, O., Ganaiem, M., Sragovich, S., Lobyntseva, A., Giladi, E., Yeheskel, A., Avitan, A., Vatine, G. D., Gurwitz, D., Ivashko-Pachima, Y., & Gozes, I. (2021). Introducing ADNP and SIRT1 as new partners regulating microtubules and histone methylation. Molecular Psychiatry, 26(11), 6550–6561. 10.1038/s41380-021-01143-9

Ivashko-Pachima, Y., & Gozes, I. (2020). Activity-dependent neuroprotective protein (ADNP)-end-binding protein (EB) interactions regulate microtubule dynamics toward protection against tauopathy. Progress in Molecular Biology and Translational Science, 177, 65–90. 10.1016/bs.pmbts.2020.07.008

Kapitansky, O., Karmon, G., Sragovich, S., Hadar, A., Shahoha, M., Jaljuli, I., Bikovski, L., Giladi, E., Palovics, R., Iram, T., & Gozes, I. (2020). Single cell ADNP Predictive of human muscle disorders: Mouse knockdown results in muscle wasting. Cells, 9(10), 2320. 10.3390/cells9102320

Karmon, G., Sragovich, S., Hacohen-Kleiman, G., Ben-Horin-Hazak, I., Kasparek, P., Schuster, B., Sedlacek, R., Pasmanik-Chor, M., Theotokis, P., Touloumi, O., Zoidou, S., Huang, L., Wu, P. Y., Shi, R., Kapitansky, O., Lobyntseva, A., Giladi, E., Shapira, G., Shomron, N., … Gozes, I. (2021). Novel ADNP syndrome mice reveal dramatic Sex-Specific peripheral gene expression with brain synaptic and TAU pathologies. Biological Psychiatry, 92(1), 81–95. 10.1016/j.biopsych.2021.09.018

Kitajima, Y., & Ono, Y. (2016). Estrogens maintain skeletal muscle and satellite cell functions. Journal of Endocrinology, 229(3), 267–275. 10.1530/joe-15-0476

NHS. (2023, July 21). Who can and cannot take continuous combined HRT. nhs.uk. https://www.nhs.uk/medicines/hormone-replacementtherapy-hrt/continuous-combined-hormonereplacement-therapy-hrt-tablets-capsules-and-patches/who-can-and-cannot-take-continuouscombined-hrt/

NHS. (2023, September 22). Side effects of hormone replacement therapy (HRT). nhs.uk. https://www.nhs.uk/medicines/hormone-replacement-therapy-hrt/side-effects-of-hormonereplacement-therapy-hrt/

Oseni, T., Patel, R., Pyle, J., & Jordan, V. (2008). Selective estrogen receptor modulators and phytoestrogens. Planta Medica, 74(13), 1656–1665. 10.1055/s-0028-1088304

Pinhasov, A., Mandel, S., Torchinsky, A., Giladi, E., Pittel, Z., Goldsweig, A. M., Servoss, S. J., Brenneman, D. E., & Gozes, I. (2003). Activity-dependent neuroprotective protein: a novel gene essential for brain formation. Developmental Brain Research, 144(1), 83–90. 10.1016/s0165-3806(03)00162-7

Russell, J. K., Jones, C. K., & Newhouse, P. A. (2019). The role of estrogen in brain and cognitive aging. Neurotherapeutics, 16(3), 649–665. 10.1007/s13311-019-00766-9

Sex hormones and health. (2017, February 1). PubMed. https://pubmed.ncbi.nlm.nih.gov/30633468/

Shapira, G., Karmon, G., Hacohen-Kleiman, G., Ganaiem, M., Shazman, S., Theotokis, P., Grigoriadis, N., Shomron, N., & Gozes, I. (2024). ADNP is essential for sex-dependent hippocampal neurogenesis, through male unfolded protein response and female mitochondrial gene regulation. Molecular Psychiatry, 30(6), 2696–2706. 10.1038/s41380-024-02879-w

Souza, V. R., Mendes, E., Casaro, M., Antiorio, A. T. F. B., Oliveira, F. A., & Ferreira, C. M. (2018). Description of ovariectomy protocol in mice. Methods in Molecular Biology, 1916, 303–309. 10.1007/978-1-4939-8994-2_29

Tao, X., Yan, M., Wang, L., Zhou, Y., Wang, Z., Xia, T., Liu, X., Pan, R., & Chang, Q. (2020). Effects of estrogen deprivation on memory and expression of related proteins in ovariectomized mice. Annals of Translational Medicine, 8(6), 356. 10.21037/atm.2020.02.57

Vulih-Shultzman, I., Pinhasov, A., Mandel, S., Grigoriadis, N., Touloumi, O., Pittel, Z., & Gozes, I. (2007). Activity-Dependent Neuroprotective protein snippet NAP reduces TAU hyperphosphorylation and enhances learning in a novel transgenic mouse model. Journal of Pharmacology and Experimental Therapeutics, 323(2), 438–449. 10.1124/jpet.107.129551

Xu, Z., Yu, Z., Chen, M., Zhang, M., Chen, R., Yu, H., Lin, Y., Wang, D., Li, S., Huang, L., Li, Y., Yuan, J., & Yin, P. (2022). Mechanisms of estrogen deficiency-induced osteoporosis based on transcriptome and DNA methylation. Frontiers in Cell and Developmental Biology, 10, 1011725. 10.3389/fcell.2022.1011725

